# Monoclonal antibodies targeting the influenza virus N6 neuraminidase

**DOI:** 10.1101/2021.02.15.431355

**Authors:** Shirin Strohmeier, Fatima Amanat, Juan Manuel Carreño, Florian Krammer

**Author notes:** Address correspondence to Florian Krammer.

## Abstract

Influenza A viruses are a diverse species that include 16 hemagglutinin (HA) subtypes and 9 neuraminidase (NA) subtypes. While the antigenicity of many HA subtypes is reasonably well studied, less is known about NA antigenicity, especially when it comes to non-human subtypes that only circulate in animal reservoirs. The N6 NA subtypes are mostly found in viruses infecting birds. However, they have also been identified in viruses that infect mammals, such as swine and seals. More recently, highly pathogenic H5N6 subtype viruses have caused rare infections and mortality in humans. Here, we generated murine mAbs to the N6 NA, characterized their breadth and antiviral properties *in vitro* and *in vivo* and mapped their epitopes by generating escape mutant viruses. We found that the antibodies had broad reactivity across the American and Eurasian N6 lineages, but relatively little binding and inhibition of the H5N6 NA. Several of the antibodies exhibited strong NA inhibition activity and some also showed activity in the antibody dependent cellular cytotoxicity reporter assay and neutralization assay. In addition, we generated escape mutant viruses for six monoclonal antibodies and found mutations on the lateral ridge of the NA. Lastly, we observed variable protection in H4N6 and H5N6 mouse challenge models when the antibodies were given prophylactically.

**Importance:** The N6 NA has recently gained prominence due to the emergence of highly pathogenic H5N6 viruses. Currently, there is limited characterization of the antigenicity of avian N6 neuraminidase. Our data is an important first step towards a better understanding of the N6 NA antigenicity.

## Introduction

Influenza A viruses are categorized based on 16 hemagglutinin (HAs) subtypes and 9 neuraminidase (NAs) subtypes which are all found in the avian reservoirs (1). Two HA-like and NA-like virus isolates have also been found in bats (2, 3). HAs facilitate binding and entry into host cells and immune responses to HA have been correlated with protection from infection and/or disease. As such, the antigenicity of numerous HAs is highly characterized. Despite increased efforts to better understand NA antigenicity of human seasonal N1, N2 and type B NAs (4–14) for the design of improved influenza virus vaccines, little work has been done on non-human NA subtypes (group 1: N1, N4, N5, N8; group 2: N2, N3, N6, N7, N9), and most work has focused almost exclusively on the N9 NA (15–20).

Viruses composed of the N6 NA circulate in avian species in combination with all 16 HA subtypes, although H3N6 (21), H4N6 (22), H5N6 (22), H6N6 (23) and H13N6 (24) are most common. In addition, H3N6, H4N6, H5N6 and H6N6 have been isolated from swine (25–28), H4N6 viruses have been found in marine mammals (29) and H5N6 viruses have been found in cats (22). Avian H7N6 viruses have also been shown to be transmissible in guinea pigs (30). The N6 subtype can be further divided into two lineages, a predominantly Eurasian lineage (EAL) and a predominantly North American lineage (NAL) (**Figure 1**) (1), although isolates from North America sometimes fall into the Eurasian lineage and *vice versa.* Some N6 isolates have also been found to be resistant to NA inhibitors, which is a concern (31, 32). Another peculiarity that seems to be specific for N6 NAs is the ability of some strains to aid cleavage of HA in concert with thrombin-like proteases, as observed for an H7N6 virus isolate (33), which allows systemic replication of these viruses without the presence of a polybasic cleavage site of the HA.

**Figure 1.**
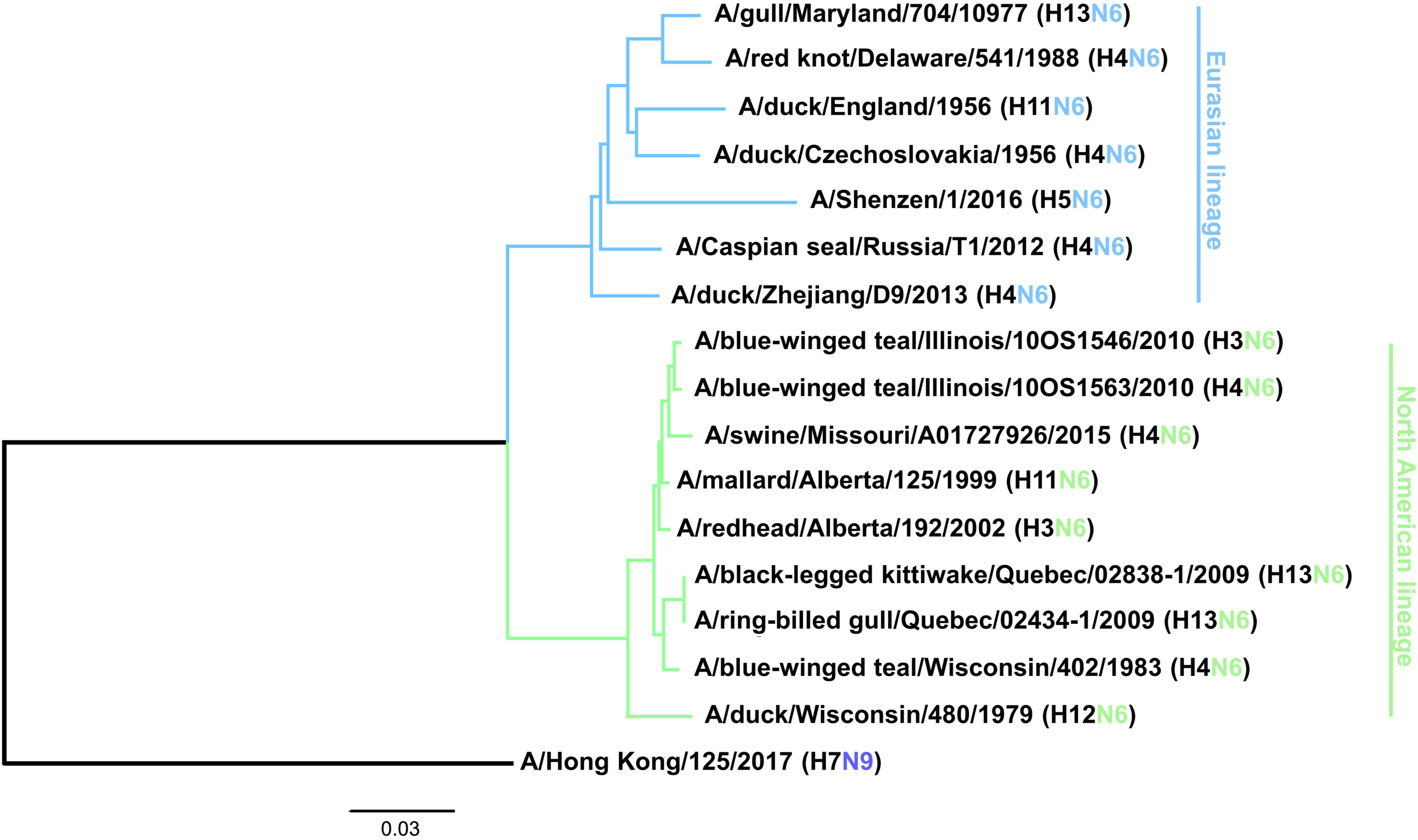
Phylogenetic tree of the N6 subtype based on amino acid sequences. A phylogenetic tree was constructed based on the amino acid sequence alignment of the various N6 NAs. N9 was used as an outgroup and the scale bar represents a 3% change in amino acid sequence. Eurasian and North American lineages are indicated. The tree was constructed in Clustal Omega and visualized in FigTree.

With the emergence of H5NX viruses in the early 2000s (34), highly pathogenic H5N6 viruses hosting a Eurasian N6 appeared in chickens in 2013 and have since 2015 also infected humans and have - according to the World Health Organization (WHO) as of December 18^th^ 2020 (35) - led to 8 fatalities (36–40). Therefore, H5N6 vaccine candidates are being prepared (e.g. (H5N6)-PR8-IDCDC-RG42A). While these vaccines are focused on inducing anti-HA antibodies, it would also be important to better understand the antigenicity of the N6 NA. Here, using murine hybridoma technology, we generated and characterized monoclonal antibodies (mAbs) to better understand breadth, functionality, epitopes and protective effect of the antibody response to N6 NA.

## Results

### Generation of mAbs

To generate and isolate mAbs, we vaccinated mice first with A/swine/Missouri/A01727926/2015 (H4N6, NAL) twice in a three-week interval (41). Three weeks later mice were immunized with A/Shenzhen/1/2016 (H5N6, EAL). After another three weeks, one animal was boosted with A/duck/Zhejiang/D9/2013 (H4N6, EAL), three days later the animal was euthanized, the spleen was removed and a hybridoma fusion performed. Ten mAbs with strong reactivity to a recombinant version of the N6 of A/swine/Missouri/A01727926/2015 were identified as a result of the hybridoma fusion (**Table 1**).

**Table 1:** Monoclonal antibodies characterized in this study and their IgG subtypes

| Anti-N6 mAb | Subtype |
| --- | --- |
| KL-N6-1A5 | IgG2a |
| KL-N6-1D4 | IgG2a |
| KL-N6-1G9 | IgG2a |
| KL-N6-2E8 | IgG2a |
| KL-N6-3B5 | IgG2a |
| KL-N6-3C6 | IgG2a |
| KL-N6-3C7 | IgG2a |
| KL-N6-3D5 | IgG2a |
| KL-N6-3E1 | IgG2a |
| KL-N6-3F10 | IgG2a |

### Characterization of antibody binding breadth

First, we wanted to characterize the binding breadth of the ten mAbs. To do this, we recombinantly expressed the N6 of A/swine/Missouri/A01727926/2015 (H4N6, NAL), A/Shenzhen/1/2016 (H5N6, EAL), A/Caspian seal/Russia/T1/2012 (H4N6, EAL) and A/duck/Zhejiang/D9/2013 (H4N6, EAL) as soluble tetramers. These proteins were used as substrates for enzyme-linked immunosorbent assays (ELISAs). The N9 protein of A/Anhui/1/2013 (H7N9) was used as a negative control to ensure that mAbs were specific to the N6. All mAbs, except KL-N6-3F10, displayed strong reactivity to the N6 NA of A/swine/Missouri/A01727926/2015 (NAL, **Figure 2A**). MAb KL-N6-3F10 while still binding, had a lower endpoint titer compared to the other nine mAbs. Binding to the N6 NA of A/duck/Zhejiang/D9/2013 (EAL) and A/Caspian seal/Russia/T1/2012 (EAL) was more variable, but all mAbs reacted well in an ELISA assay and better than the positive control (**Figure 2B and C**). Interestingly, binding to the recombinant N6 NA of A/Shenzhen/1/2016 (EAL) was low for most mAbs, and only mAb KL-N6-3F10 showed decent binding. Lower binding activity was observed for KL-N6-3B5, KL-N6-3C5 and KL-N6-3E1 (**Figure 2D**). None of the mAbs bound to recombinant N9 NA which shows that antibodies were specific to the N6 (**Figure 2E**). In addition to the ELISA, we also performed an immunofluorescence assay (IF) of Madin Darby canine kidney (MDCK) cells infected with wild type isolates and re-assortant viruses from both lineages. In this assay, a majority of mAbs bound to all six tested EAL viruses (**Figure 3**). Interestingly, more antibodies showed binding to A/Shenzhen/1/2016 including KL-N6-2E8, KL-N6-3B5, KL-N6-3C5, KL-N6-3C7, KL-N6-3D5 KL-N6-3E1 and KL-N6-3F10 in the IF assay. Similar to the EAL lineage, all mAbs bound to cells infected by all eight NAL viruses in this assay (**Figure 4**).

**Figure 2.**
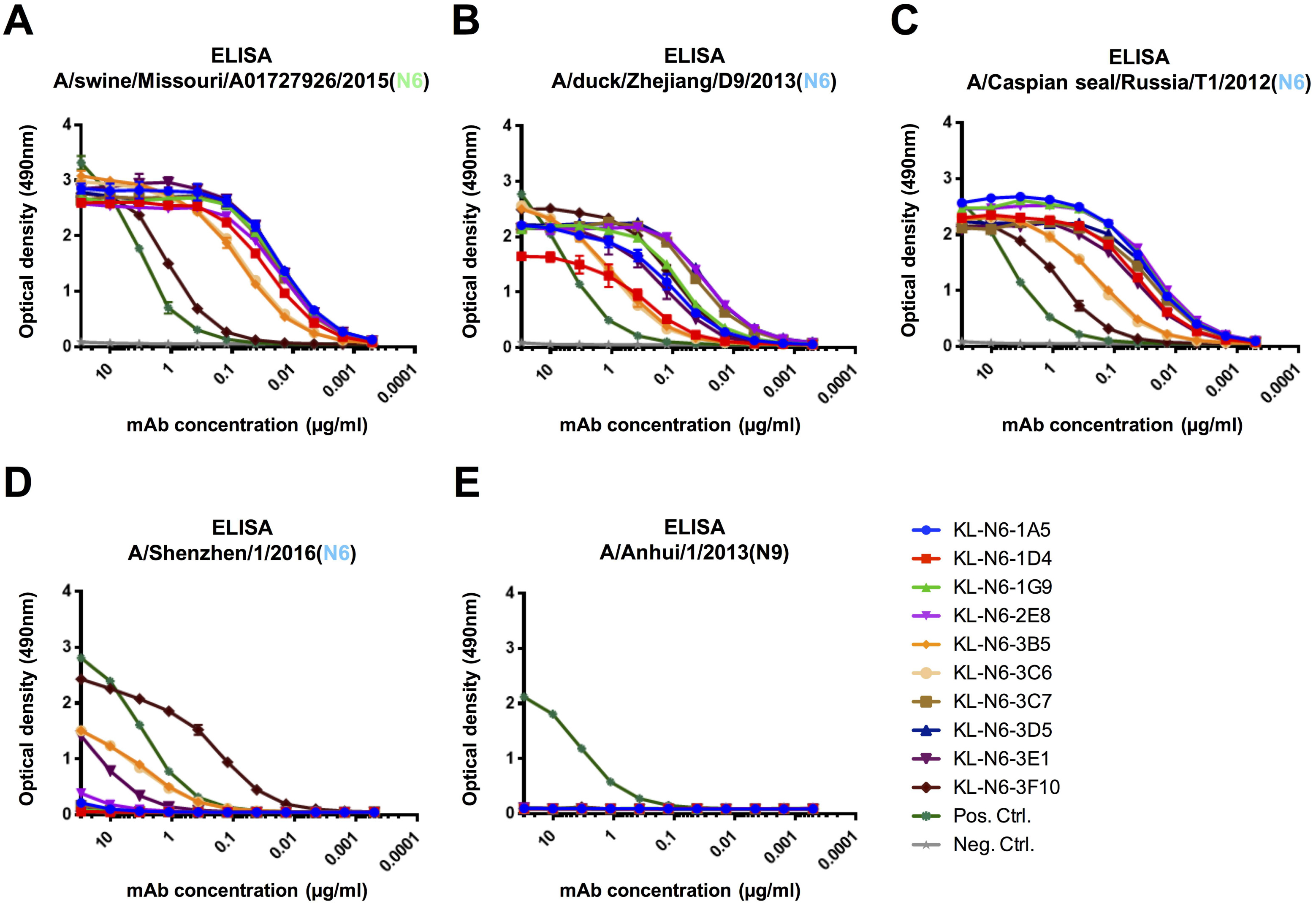
Murine N6 NA mAbs are highly specific to the N6 subtype and show broad cross-reactivity to both Eurasian and North American lineage NAs via ELISA. Binding of murine N6 NA mAbs against A/swine/Missouri/A01727926/2015 (A), A/duck/Zhejiang/D9/2013 (B), A/Caspian seal/Russia/T1/2012 (C) and A/Shenzhen/1/2016 (D) N6 NA recombinant proteins. Binding of murine N6 NA mAbs against A/Anhui/1/2013 (E) N9 NA recombinant protein. None of the antibodies showed reactivity against recombinant N9 protein, which is the closest related NA to N6. An anti-his-tag antibody was used as a positive control and an irrelevant anti-Lassa antibody (KL-AV-1A12) was used as a negative control.

**Figure 3.**
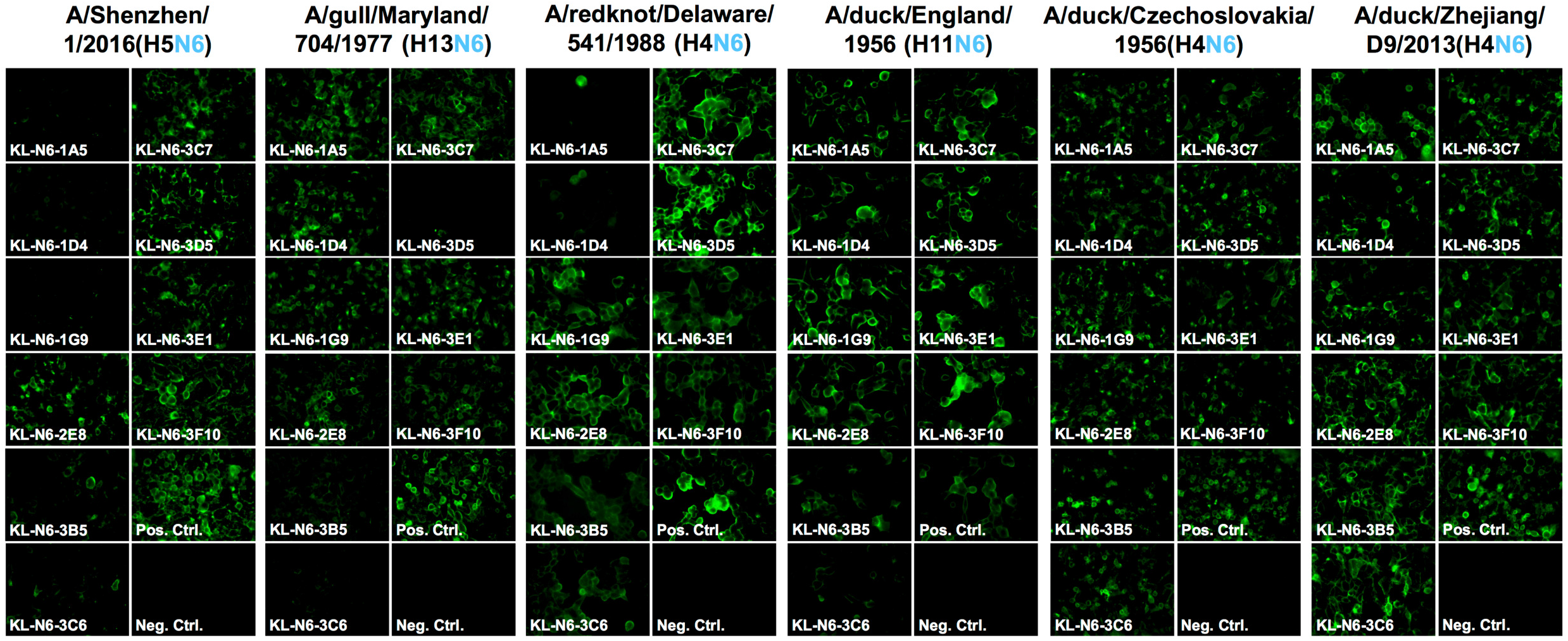
Murine N6 NA mAb binding to Eurasian lineage virus infected MDCK cells. MDCK cells were infected at a multiplicity of infection (MOI) of 3 with Eurasian lineage viruses and incubated for 16h. The infected cells were then fixed and the N6 mAbs were added at a concentration of 30 ug/mL. Most mAbs showed broad cross reactivity, except against A/Shenzhen/1/2016, were only 8 out of 10 mAbs show binding. MAb CR9114 was used as positive control and an anti-Lassa antibody KL-AV-1A12 as negative control.

**Figure 4.**
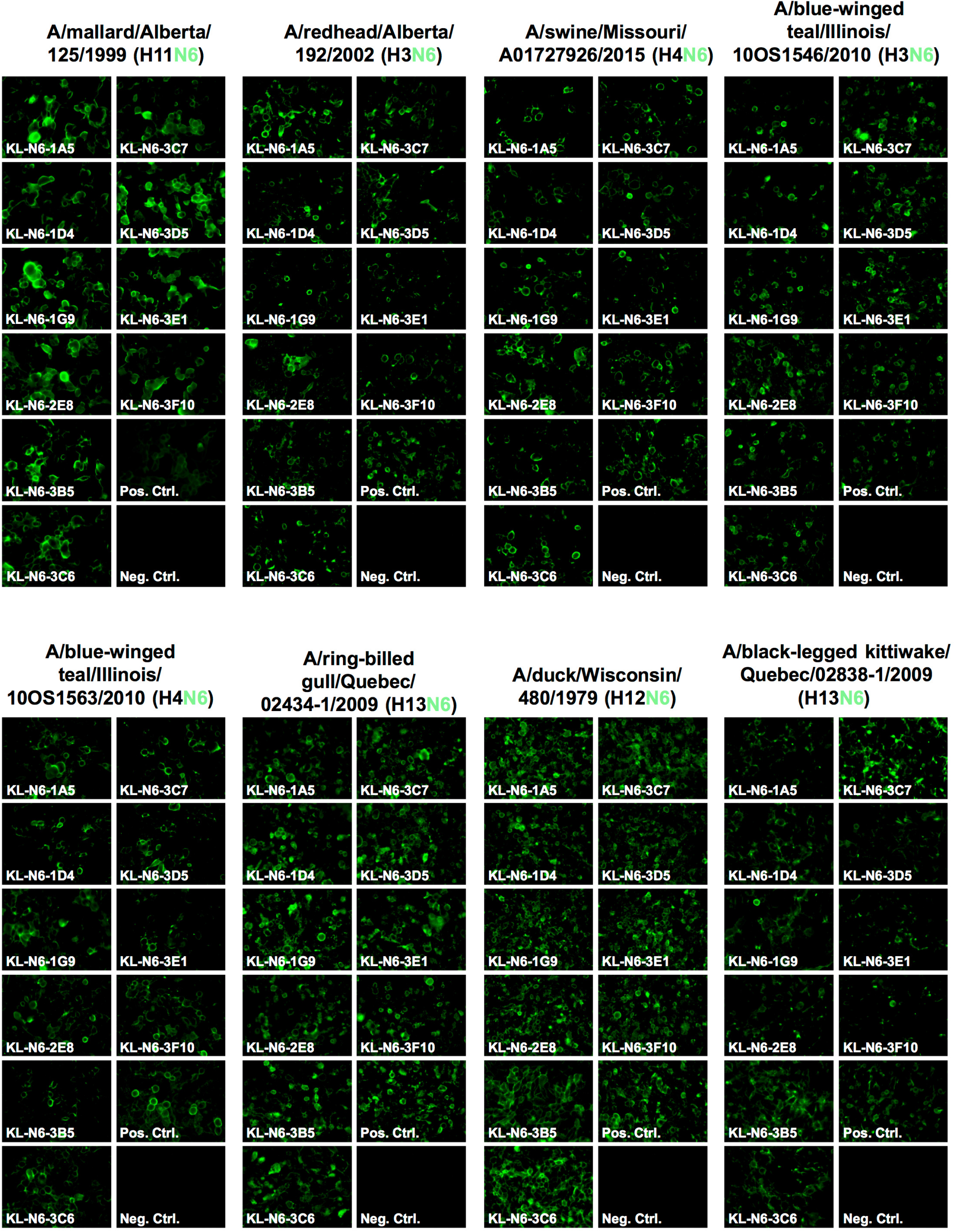
Murine N6 NA mAb binding of North American lineage virus infected MDCK cells. MDCK cells were infected at a multiplicity of infection of 3 with North American lineage viruses and incubated for 16h. The infected cells were then fixed and the N6 mAbs were added at a concentration of 30 ug/mL. The N6 mAbs show broad cross reactivity within the subset of North American lineage viruses. MAb CR9114 was used as positive control and an anti-Lassa antibody KL-AV-1A12 as negative control.

### *In vitro* functional characterization of mAbs

Next, we wanted to test the *in vitro* functionality of the isolated mAbs in NA inhibition (NI), plaque reduction neutralization and antibody-dependent cellular cytotoxicity (ADCC) reporter assays. NI was measured against A/duck/Czechoslovakia/1956 (EAL, H4N6), A/swine/Missouri/A01727926/2015 (NAL, H4N6) and two versions of A/Shenzhen/1/2016 (EAL, H5N6). MAbs KL-N6-2E8, KL-N6-3C7 and KL-N6-3D5 showed the strongest inhibition of A/duck/Czechoslovakia/1956 followed by KL-N6-1G9, KL-N6-1A5, KL-N6-3F10, KL-N6-1D4, KL-N6-3E1, KL-N6-3B5 and finally KL-N6-3C6 (**Figure 5A**). The order was similar for A/swine/Missouri/A01727926/2015 except that KL-N6-3F10 was only slightly better than KL-N6-3B5 and finally KL-N6-3C6 (**Figure 5B**). NI activity was very weak for most mAbs against A/Shenzhen/1/2016, and only KL-N6-3F10 showed reasonable activity, albeit even that was low (**Figure 5C**). Of note, HA stalk-reactive antibody CR9114 (42, 43), which can inhibit NA through steric hindrance (44), was used as positive control. CR9114 performed relatively well against A/Shenzhen/1/2016. The N6 NA of A/Shenzhen/1/2016 has a stalk deletion, which is known to confer pathogenicity to highly pathogenic avian influenza (HPAI) viruses and we reasoned that a longer stalk may make the NA more vulnerable to NI. We, therefore, created a loss of function version of A/Shenzhen/1/2016 which featured a regular stalk length. While no improvement was seen for the other mAbs, KL-N6-3F10 performed much better against the ‘long stalk’ version of A/Shenzhen/1/2016 (**Figure 5D**). CR9114 performed less well, which makes sense since a longer stalk of the NA probably overcomes some of the steric hindrance exerted by anti-stalk mAbs.

**Figure 5.**
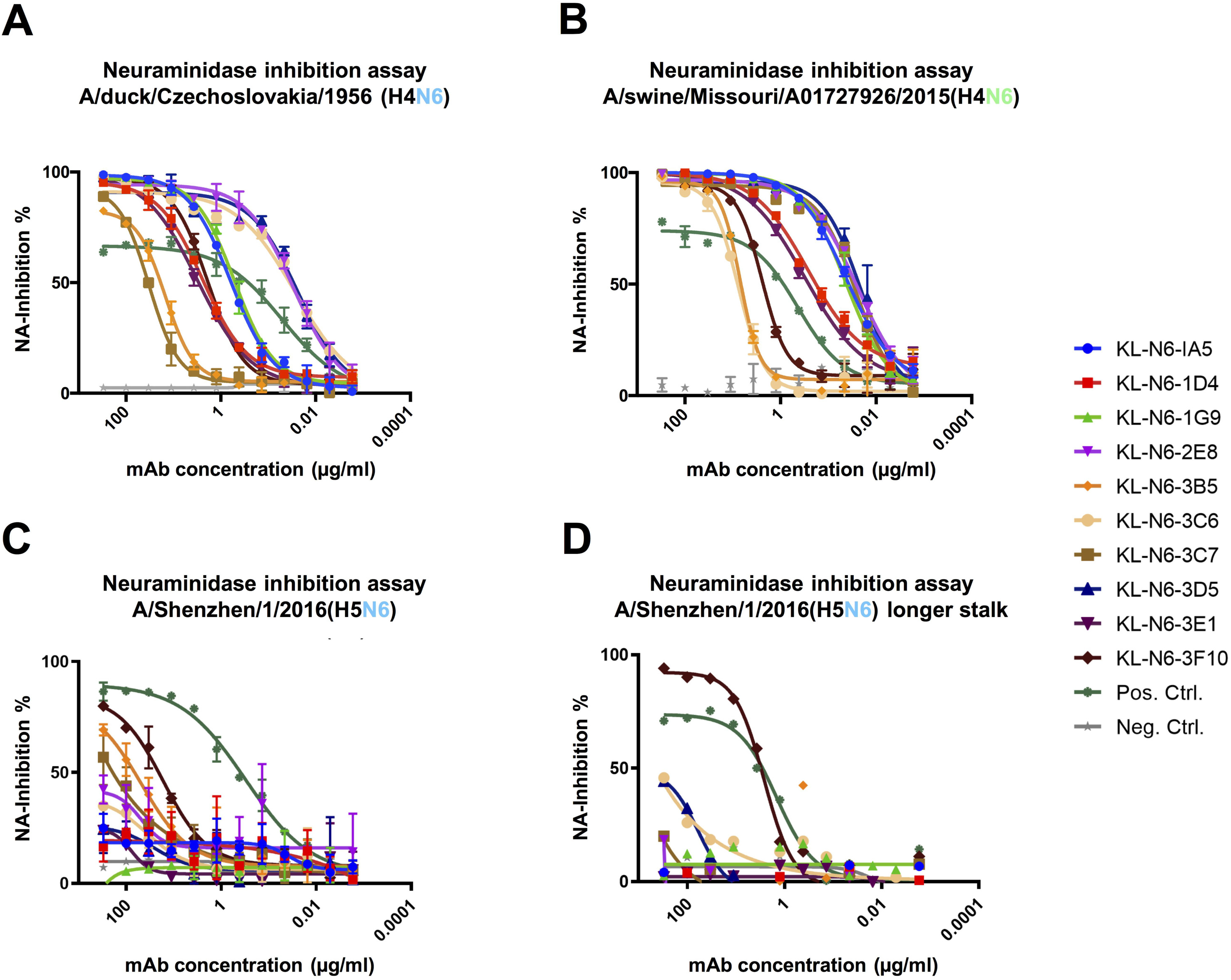
N6 mAbs strongly inhibit NA enzymatic activity. N6 NA mAbs were tested for inhibition of A/duck/Czechoslovakia/1956 (H4N6) (A), A/swine/Missouri/A01727926/2015 (H4N6) (B), A/Shenzhen/1/2016 (H5N6, low path 6:2 A/PR/8/34 reassortant lacking the polybasic cleavage site) (C) and A/Shenzhen/1/2016 with an extended N6 stalk domain (D) in the NA inhibition assay. The broad HA stalk antibody CR9114 was used as a positive control (steric hindrance-based NA inhibition) and an irrelevant anti-Lassa antibody KL-AV-1A12 was used as negative control.

Using A/duck/Czechoslovakia/1956 (H4N6), we also performed plaque reduction neutralization assays (PRNAs) to measure both reduction of plaque size (a common feature of anti-NA mAbs) and reduction of plaque numbers (usually referred to as neutralization). All mAbs reduced plaque size to various degrees, mostly in accordance with their NI activity (**Figure 6A**). Plaque number was only reduced effectively by mAbs KL-N6-1A5, KL-N6-1D4, KL-N6-1G9, KL-N6-3B5, KL-N6-3D5 and KL-N6-3E1 (**Figure 6B**). Interestingly, these results are not in concordance with the NI activity suggesting that neutralization requires a different mechanism other than NI activity for these mAbs.

**Figure 6.**
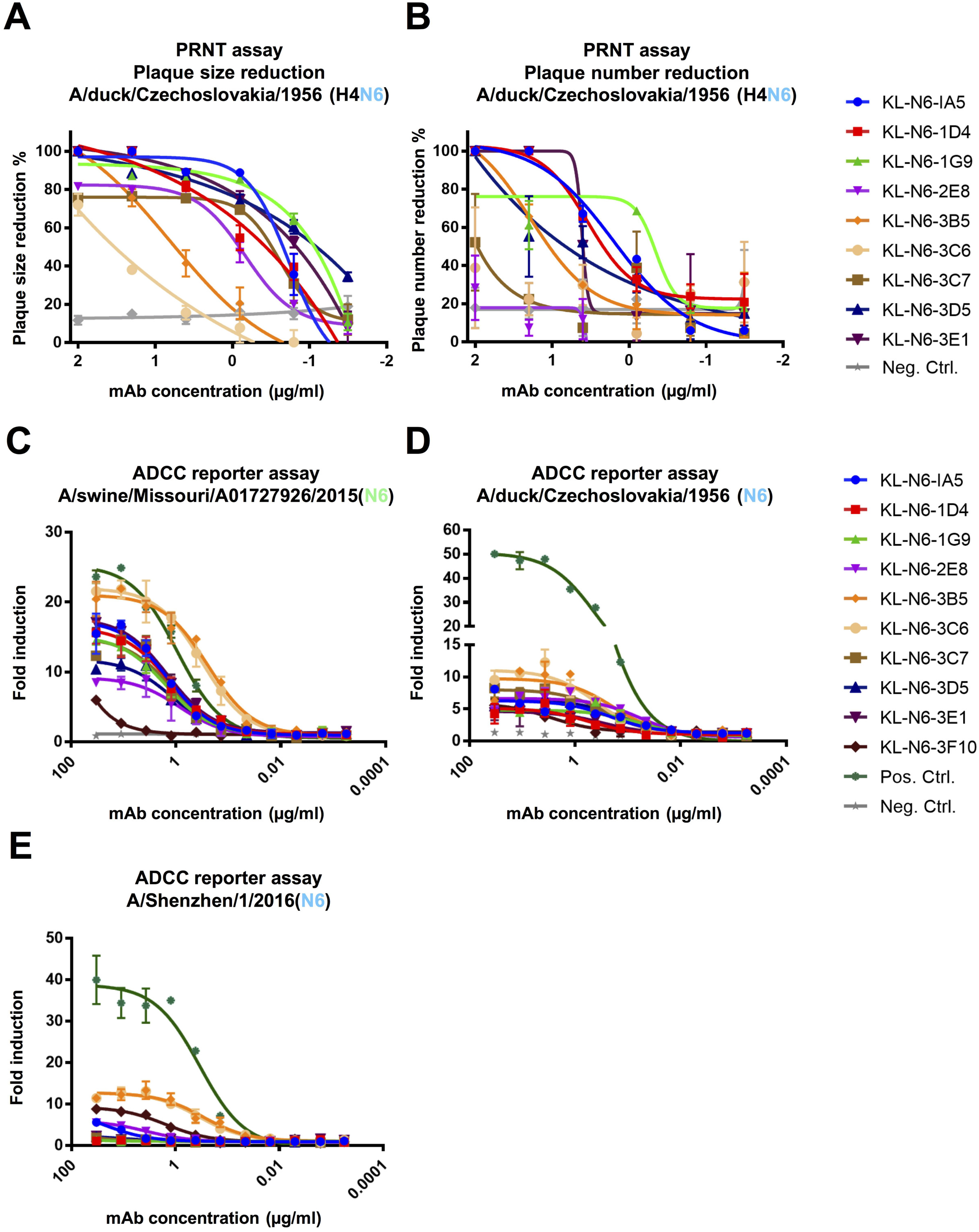
*In vitro* functional characterization of the N6 mAbs. (A) Plaque size and (B) number reduction measured via a plaque reduction neutralization assay. (C-E) ADCC reporter activity of mAbs using infected MDCK cells as the substrate. Luminescence was measured at the end of the assay to assess ADCC activity. The HA stalk antibody CR9114 was used as a positive control and an irrelevant anti-Lassa antibody KL-AV-1A12 as negative control.

ADCC reporter activity was also measured since it has been shown to be important for the protective effect of some NA mAbs. All antibodies except KL-N6-3F10 exhibited reasonable but variable ADCC reporter activity against A/swine/Missouri/A01727926/2015. Especially KL-N6-3B5 and KL-N6-3C6 showed outstanding activity that was higher than that of the positive control, stalk mAbs CR9114 which is known for its ADCC activity (**Figure 6C**). Interestingly, the ADCC activity of the mAbs was much lower against cells infected with A/duck/Czechoslovakia/1956 (**Figure 6D**) or A/Shenzhen/1/2016 (**Figure 6E**), but also here KL-N6-3B5 and KL-N6-3C6 performed better than the other mAbs.

### Epitope mapping via escape mutant generation

To learn more about the epitopes of the generated mAbs, we generated escape mutant viruses (EMVs) by subjecting A/duck/Czechoslovakia/1956 to increasing antibody pressure by passaging the virus continually with increasing amounts of antibody. An irrelevant mAb directed against the Lassa virus glycoprotein was included as control (45). We succeeded in generating six different EMVs. The EMV selected by KL-N6-1A5 showed a strong reduction in NI compared to wild type. This EMV carried a G221E mutation on the lateral side of the NA (**Figure 7**). Similar phenotypes were seen for the KL-N6-2E8 (N249S) and KL-N6-3E1 (G221E) EMVs. For the KL-N6-1D4 (R250G), KL-N6-1G9 (R250K) and KL-N6-3D5 (P246Q) EMVs, a complete loss of NI was observed. When the six mAbs were then cross-tested with the six EMVs, we found that the KL-N6-1A5 EMV (G221E) also strongly reduced NI of KL-N6-1D4 and KL-N6-1G9 while KL-N6-2E8 and KL-N6-3D5 still showed good NI (**Figure 8**). The KL-N6-1D4 (R250G) mutation abolished NI of all mAbs to a large degree. The KL-N6-1G9 EMV (R250K) had a lower impact and only caused loss of inhibition by KL-N6-1A5 and KL-N6-1D4 in addition to KL-N6-1G9, likely owing this to its conserved nature. For the KL-N6-2E8 EMV (N249S) all mAbs retained some NI activity. The KL-N6-3D5 mutation (P249Q)) had an impact on KL-N6-2E8 and itself, but less so on the other mAbs. Finally, the KL-N6-3E1 EMV, which carries the same G221E mutation in the NA as the KL-N6-1A5 EMV, showed a similar pattern as the KL-N6-1A5 EMV. However, the impact on NI by KL-N6-3E1 was stronger, potentially due to additional mutations in non-NA genes of this EMV which could have contributed to KL-N6-3E1 specific escape. The interdependence of the EMV/mAb combinations makes sense as all escape mutations map to the same patch on the NA (**Figure 9**).

**Figure 7.**
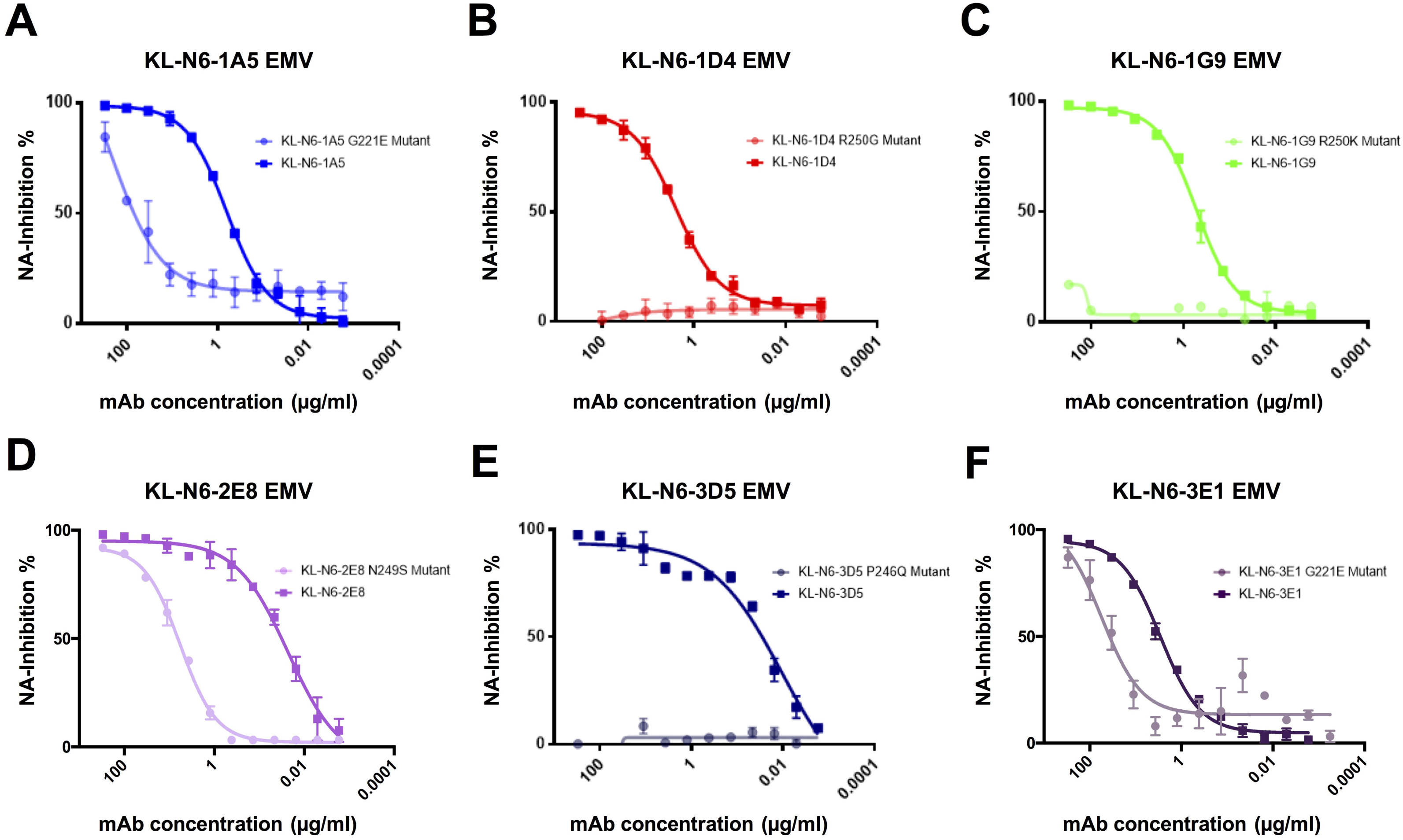
MAbs show diminished or knocked out NA inhibition against escape mutants. (A,D and F) Antibodies KL-N6-1A5, KL-N6-2E8 and KL-N6-3E1 show decreased neuraminidase inhibition activity to the respective escape mutant viruses (EMVs). EMV 1A5 and EMV 3E1 share the same mutation at position 221 (G221E), whereas EMV 2E8 has a mutation at position 249 (N249S). (B,C,E) KL-N6-1D4, KL-N6-1G9 and KL-N6-3D5 display a complete loss in neuraminidase inhibition activity against the respective EMVs. EMV 1D4 and EMV 1G9 share a mutation at the same position at aa 250 (R250G, R250K). EMV 3D5 has a mutation at position 246 (P246Q).

**Figure 8.**
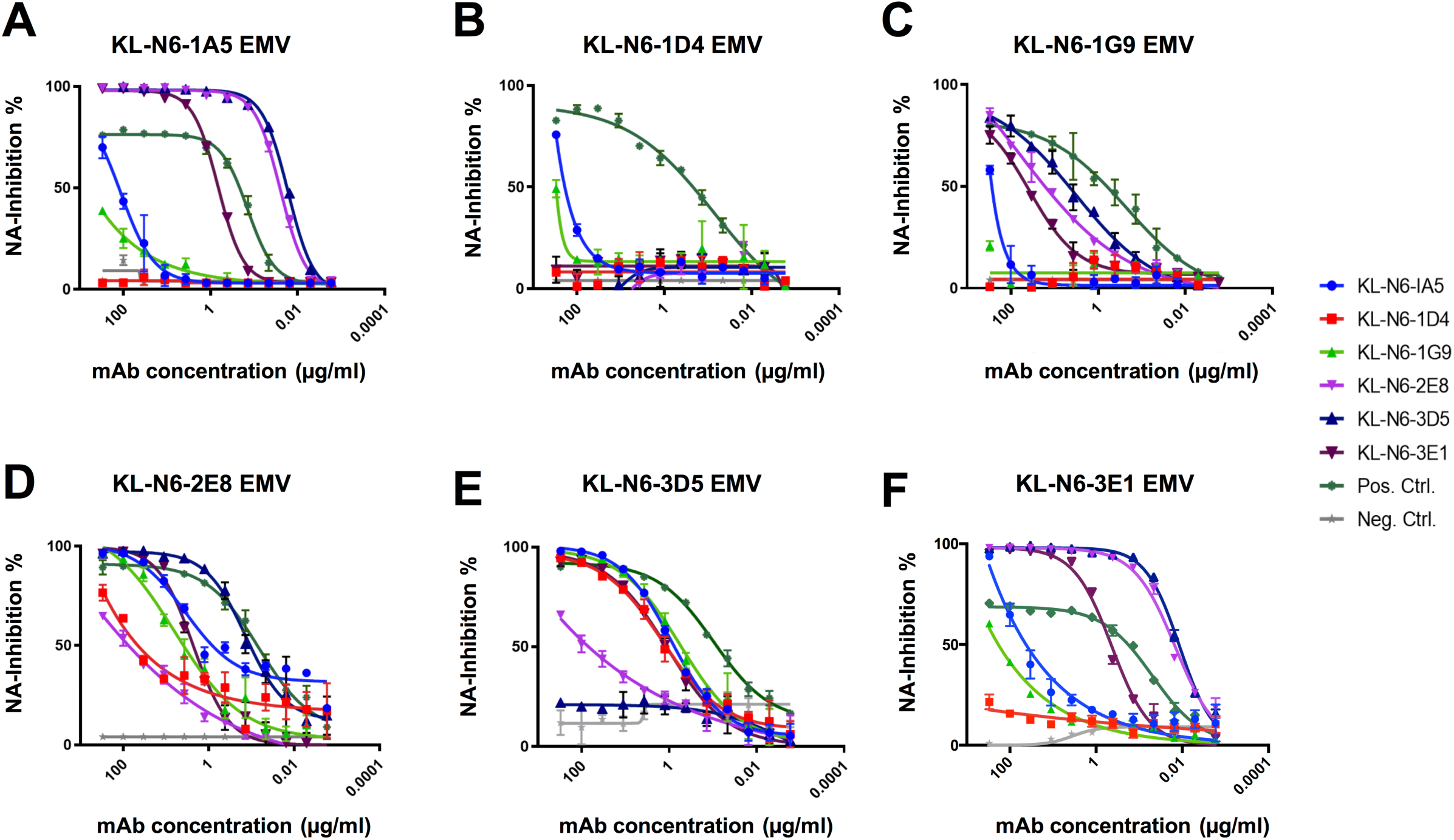
Cross-mAb NA inhibition testing of EMVs. (A) 1A5 EMV (G221E) strongly reduces NI of KL-N6-1D4 and KL-N6-1G9, whereas the inhibition capability of KL-N6-2E8 and KL-N6-3D5 fully remains. (B) 1D4 EMV (R250G) abolishes NI of all mAbs to a large degree. (C) 1G9 EMV (R250K) causes total loss of inhibition by KL-N6-1A5 and KL-N6-1D4. (D) 2E8 EMV (N249S) does not strongly impact NI inhibition. (E) 3D5 EMV (P249Q) leads to a decreased NI by KL-N6-2E8 but has lass impact on the remaining mAbs. (F) 3E1 EMV (G221) shows a similar pattern as 1A5 EMV. As a positive control the broad HA stalk antibody CR9114 was used. The negative control was anti-Lassa GP mAb KL-AV-1A12.

**Figure 9.**
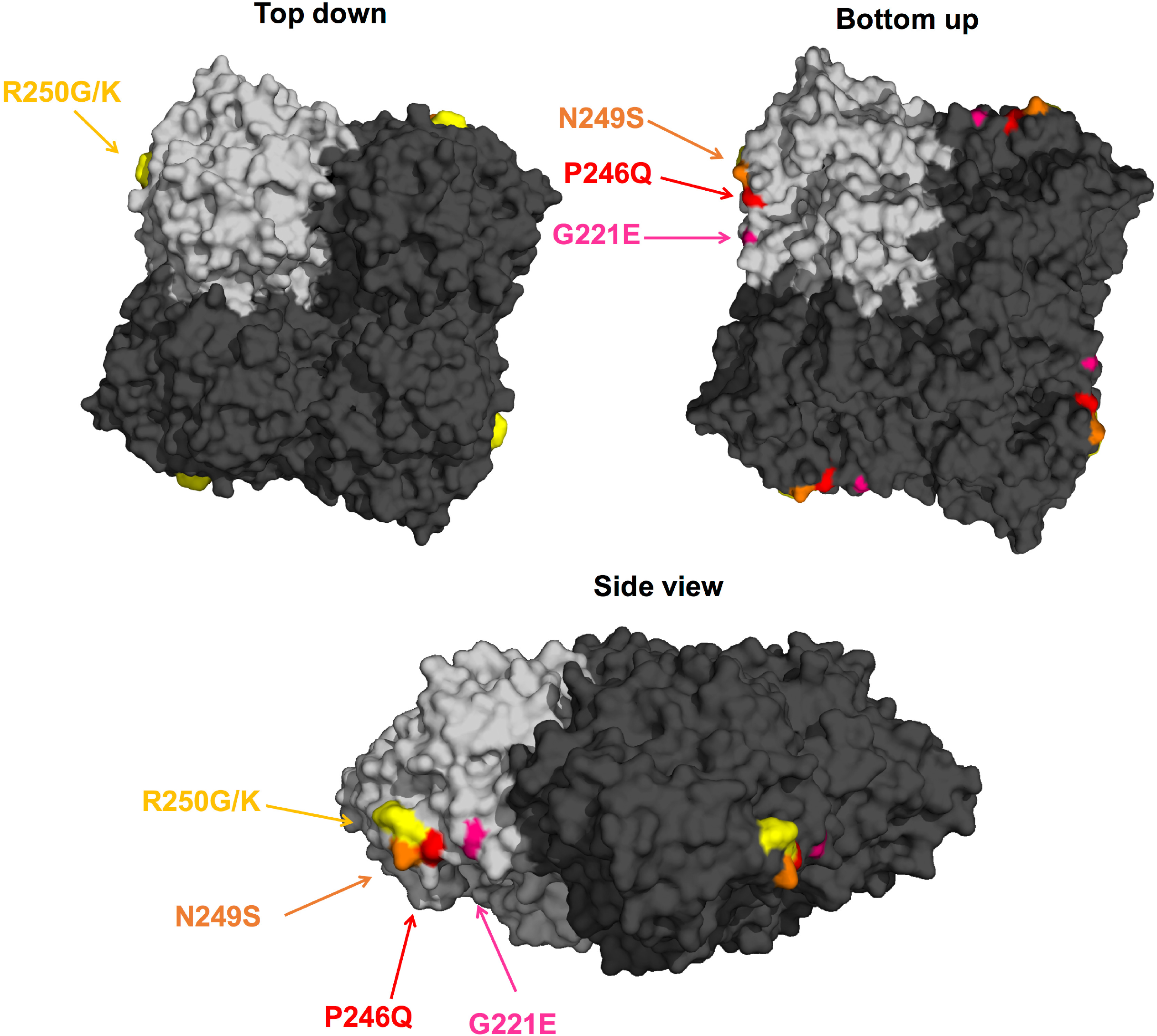
Visualization of N6 NA mAb escape mutations. The tetrameric protein of A/duck/England/1/1956 (H4N6, PDB 1V0Z) is shown in top down, bottom up and side view, one of the monomers is highlighted in light grey. The individual mutations are highlighted in different colours and are in close proximity on the lateral surface of the NA.

### *In vivo* protective effect of mAbs

Lastly, we wanted to determine the protective effect of the isolated mAbs against viral challenge in a mouse model. DBA/2J mice were administered antibody via intraperitoneal route two hours before infection with 5 mg/kg of mAb and then challenged intranasally with 5 times the 50% lethal dose (mLD_50_) of either A/duck/Czechoslovakia/1956 (H4N6, EAL) or A/Shenzhen/1/2016 (H5N6, EAL). Weight loss and survival was then monitored for 14 days. Protection from A/duck/Czechoslovakia/1956 was variable, with almost no weight loss and no mortality observed with KL-N6-1A5, KL-N6-3C6 and KL-N6-3F10 (**Figure 10A and B**). KL-N6-1D4, KL-N6-1G9 and KL-N6-3B5 were also protective but mice experienced a transient body weight loss (approximately 10%). KL-N6-3C7, KL-N6-3D5 and KL-N6-3E1 were less protective against weight loss and for KL-N6-3C7 and KL-N6-3E1 20% and 40% of the animals succumbed to infection respectively. No survival benefit was observed for any of the mAbs in the A/Shenzhen/1/2016 challenge model (**Figure 10C and D**).

**Figure 10.**
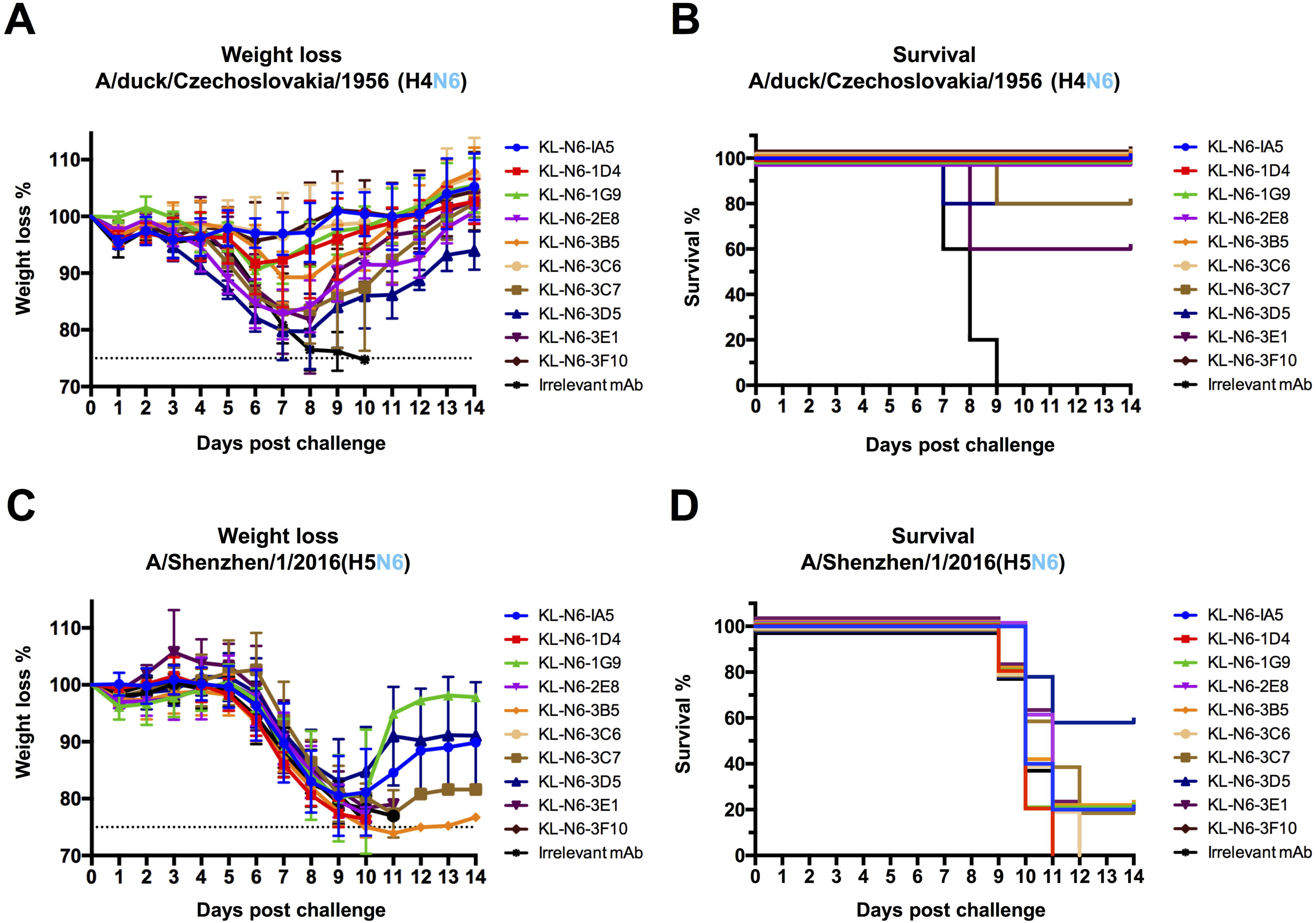
*In vivo* challenge study after prophylactic mAb treatment. N6 NA mAbs were administered to mice at a concentration of 5 mg/kg two hours prior to challenge. An irrelevant anti-Lassa antibody (KL-AV-1A12) was given as a negative control. Two hours after mAb administration, mice were challenged with 5xLD_50_ of A/duck/Czechoslovakia/1956 (H4N6) or A/Shenzhen/1/2016 (H5N6, low path 6:2 A/PR/8/34 reassortant lacking the polybasic cleavage site). Morbidity (A) and mortality (B) following A/duck/Czechoslovakia/1956 (H4N6) challenge. Morbidity (C) and mortality (D) following A/Shenzhen/1/2016 (H5N6, low path 6:2 A/PR/8/34 reassortant lacking the polybasic cleavage site) challenge.

## Discussion

In this study, we are shedding first light on antigenicity of the N6 NA. As expected, significant cross reactivity exists. This could be due to our immunization regimen which might have favoured cross reactive epitopes but it may also be a consequence of the absence of antigenic drift for most N6 NAs. The characterized mAbs also showed strong NI activity against isolates from the Eurasian and the North American lineage and protected - to various degrees - against challenge with a historic Eurasian lineage H4N6 strain.

However, binding to the N6 of a HPAI H5N6 was suboptimal. Binding to recombinant proteins was only observed for a fraction of the mAbs. Some mAbs that did not bind the recombinant protein but were able to bind cells infected with H5N6 well, suggesting suboptimal folding of the recombinant protein. Nevertheless, most of the mAbs – except KL-N6-3F10, showed negligible NA inhibition activity against H5N6. The lack of functional cross reactivity might suggest that the N6 of H5N6 has experienced more immune pressure than ‘regular’ avian isolates, potentially resulting in some antigenic drift.

Another feature of this N6 is its truncated stalk. We hypothesized that a longer stalk would maybe make the NA more susceptible to NI by the mAbs but this was only the case for KL-N6-3F10, not for other mAbs. Interestingly, CR9114, an anti-stalk mAb that can exhibit NI through steric hindrance, showed more activity to the H5N6 virus with the truncated stalk than against the version with the long stalk. This also makes sense since a longer stalk displaces the NA enzymatic site further from HA stalk bound mAbs. Disappointingly, none of the mAbs showed protection against H5N6 in a mouse model. Potential explanations could be the higher pathogenicity of H5N6 (even though a A/PR/8/34-based vaccine strain without polybasic cleavage site was used) or the mouse model which lacks complement components. This, paired with lower antibody binding and NI inhibition likely explains the failure to protect the animals. Of note, the 5 mg/kg dose, while standard in our laboratory, is relatively low. It is therefore possible that higher doses would have been protective.

We also mapped the putative epitopes of six of the mAbs through generating escape mutants. Interestingly, all escape mutations are in close proximity on the lateral surface of the NA. This patch is reminiscent of the binding side of CD6 and other N1 specific and B NA specific antibodies that may span two monomers (8, 13, 46).

While our work provides first insights into the antigenicity of the N6 NA, much more work is needed, especially given that cross reactivity to the N6 of H5N6 was limited and not protective. As such, the N6 NA needs to be further explored and potential antigenic drift mapped.

## Material and Methods

### Viruses

The viruses A/duck/Wisconsin/480/1979 (H12N6; BEI Resources # NR-28616), A/redhead/Alberta/192/2002 (H3N6; BEI Resources # NR-45145), A/duck/England/1956 (H11N6; BEI Resources # NR-21660), A/gull/Maryland/704/1977 (H13N6; BEI Resources # NR-21663), A/black-legged kittiwake/Quebec/02838-1/2009 (H13N6; BEI Resources # NR-31217), A/ring-billed gull/Quebec/02434-1/2009 (H13N6; BEI Resources # NR-31218), A/mallard/Alberta/125/1999 (H11N6; BEI Resources # NR-45187), A/blue-winged teal/Illinois/10OS1546/2010 (H3N6; BEI Resources # NR-35981) and A/blue-winged teal/Wisconsin/402/1983 (H4N6; BEI Resources # NR-48971) were obtained from the Biodefense and Emerging Infections Research Resources Repository (BEI Resources). The viruses were propagated in 10 day old embryonated chicken eggs (Charles River Laboratories) at 37°C for 2 days. The viral titers were determined by conducting a standard plaque assay on Madin-Darby Canine Kidney (MDCK) cells as described earlier (47). The viruses A/duck/Czechoslovakia/1956 (H4N6), A/Shenzhen/1/2016 (H5N6) and A/swine/Missouri/A01727926/2015 (H4N6) were rescued in the backbone of A/PR/8/34 as a 6:2 reassortant, containing the HA and NA segments of the original virus isolates (41, 48). Importantly, the H5 HA in the 6:2 A/PR/8/34 reassortant virus that was generated lacked the polybasic cleavage site.

### Cells

MDCK cells were cultivated in Dulbecco’s Modified Eagle’s Medium (complete DMEM, Gibco) containing 1% penicillin/streptomycin (100 U/ml of penicillin, 100 μg/ml streptomycin, Gibco) antibiotic solution, 10% fetal bovine serum (FBS, Gibco) and 1% hydroxyethylpiperazine ethane sulfonic acid (HEPES). SP2/0-Ag14 myeloma cells were propagated in complete DMEM supplemented with 1% L-Glutamine (Gibco).

High five cells (BTI-TN-5B1-4, *Trichoplusia ni*) were grown in serum-free Express Five media (Gibco) containing 1% L-Glutamine and 1% penicillin/streptomycin antibiotics mix. Sf9 cells (*Spodoptera frugiperda*) adapted from the cell line ATCC CRL-1711 were maintained in Trichoplusia ni medium – Fred Hink (TNM-FH, Gemini Bioproducts) supplemented with 1% penicillin/streptomycin antibiotics mix, 1% pluronic F-68 (Sigma-Aldrich) and 10% fetal bovine serum. In order to passage the baculoviruses, the media was switched to 3% TNM-FH (1% penicillin/streptomycin, 1% pluronic F-68, 3% FBS).

### Recombinant proteins

The recombinant N6 glycoproteins used in this study were expressed in insect cells using the baculovirus expression system. The globular head domain of the respective N6 protein from the strains A/duck/Czechoslovakia/1956 (H4N6), A/Shenzen/1/2016 (H5N6), A/swine/Missouri/A01727926/2015 (H4N6), A/Caspian seal/Russia/T1/2012 (H4N6) and A/duck/Zhejiang/D9/2013 (H4N6) as well as for the N9 of A/Anhui/1/2013 (H7N9) were cloned into a baculovirus shuttle vector, containing a N-terminal signal peptide, followed by a hexahistidine purification tag, a VASP (vasodilator-stimulated phosphoprotein) tetramerization domain and a thrombin cleavage site (49). The baculoviruses were passaged in Sf9 cells to reach higher titers and were then used to infect High-five cells for protein expression. After three days of infection, the soluble proteins were purified from the supernatant, as previously described (50), and were then stored at −80°C for further usage.

### Enzyme-linked immunosorbent assay (ELISA)

Ninety-six well flat bottom plates (Immulon 4 HBX plates, ThermoFisher Scientific) were coated with 50 μl/well of 2 μg/ml recombinant protein in 1x coating buffer (Seracare) at 4°C overnight. The following day, the plate was washed three times with 100 μl/well of 0.1% Tween 20 (TPBS) to remove residual coating solution and then incubated with 100 μl/well of 3% milk dissolved in TPBS for 1h at room temperature to avoid non-specific binding. The blocking solution was removed and the primary antibody added at a starting concentration of 30 μg/ml in 1% milk/TPBS. The antibodies were serially diluted 1:3 and the plates were incubated for 1 hour at room temperature (RT). An anti-

Lassa antibody (KL-AV-1A12 (45)) was included as a negative control and an antihistidine antibody as a positive control. The plate was then washed three times with 100 μl/well TPBS and an anti-mouse secondary antibody (anti-mouse IgG H&L antibody peroxidase conjugated, Rockland) diluted 1:3000 in 1% milk/TBPS was added for 1h at RT. The plate was washed three times and 100 μl/well of SigmaFast o-Phenylenediamine dihydrochloride (OPD) developing solution (Sigma Aldrich) were added. The reaction was stopped after 10 min incubation at RT with 50 μl/well of 3M hydrochloric acid (HCl). The plate was read with a Synergy H1 hybrid multimode microplate reader (BioTek) at an optical density of 490 nm. The data was analysed by using GraphPad Prism 7 software.

### Generation of monoclonal N6-antibodies

Mouse mAbs were produced using hybridoma technology, as previously described (51, 52). Briefly, one female 6-8 week old BALB/c mouse (The Jackson Laboratory) was immunized via the intraperitoneal route with 10^5^ plaque forming units (PFU)/ml of A/swine/Missouri/A01727926/2015 (H4N6) in 100 μl PBS, the same virus was used 21 days later for an intranasal infection. Three weeks later, the mouse was intranasally infected with a sublethal dose of 10^3^ PFU/ml of A/Shenzhen/1/2016 (H5N6) in a volume of 50 μl. After three weeks, the mouse received a final boost intraperitoneally with 100 μg of recombinant N6 protein of A/duck/Zhejiang/D9/2013 (H4N6) adjuvanted with 10 μg of poly (I:C) (Invivogen). Three days after the final boost, the mouse was sacrificed and the spleen removed. The spleen was washed with PBS and then flushed with serum-free DMEM (1% penicillin/streptomycin) to obtain the splenocytes. The splenocytes were fused with SP2/0-Ag14 myeloma cells in a ratio of 5:1 using pre-warmed polyethylene glycol (PEG; Sigma-Aldrich). The cells were grown on semi-solid selection & cloning medium with hypoxanthine-aminopterin-thymidine (HAT; Molecular Devices) for 10 days and were then expanded to a 96-well cell culture plate.

To obtain antibody specificity against N6, the supernatant of the individual clones was screened via ELISA against recombinant NA of A/duck/Zhejiang/D9/2013 (H4N6). Reactive clones were tested using the Pierce rapid antibody isotyping kit (Life Technologies). Only IgG heavy-chain subgroups were continuously expanded. The selected hybridoma clones were first expanded in Clonacell-HY Medium E and then constantly switched to Hybridoma SFM media (Gibco) supplemented with 1% penicillin/streptomycin. The antibodies were then purified as previously described (8).

### Immunofluorescence (IF) assay

MDCK cells were seeded in a sterile 96-well cell culture plate at a density of 25,000 cells/well using complete DMEM. The next day, cells were infected with a multiplicity of infection (MOI) of 3.0 for 16h using serum free minimal essential media (MEM, Gibco), supplemented with penicillin/streptomycin, HEPES (Gibco), glutamine (200mM L-Glutamine, Gibco) and sodium bicarbonate (sodium bicarbonate 7.5% solution, Gibco). The cells were fixed with 3.7% paraformaldehyde (PFA)/PBS for 1h at RT. The plate was the blocked for 1h with 3%milk/PBS. The antibodies were diluted to 30 μg/ml in 1% milk/PBS and incubated for 1h at RT. An anti-Lassa antibody (KL-AV-1A12 (45)) was included as a negative control and CR9114 (42), a broadly cross-reactive HA antibody, as a positive control. The plate was washed three times with 1xPBS and then incubated with a fluorescence goat anti-mouse IgG heavy plus light chain (H+L)–Alexa Fluor 488 antibody (Abcam) or an anti-human secondary (Sigma) for CR9114, which were diluted 1:1000 in 1% milk/PBS. The cells were washed with PBS and kept in PBS to avoid drying out during microscopy (Olympus IX-70).

### Antibody-dependent cellular cytotoxicity (ADCC) reporter assay

To assess potential ADCC activity of the N6 mAbs the ADCC reporter bioassay kit from Promega was used (53). MDCK cells (25,000 cells/well) were seeded in a white, flat bottom 96-well cell culture plate (Corning). The following day the cells were washed with PBS and then infected with an MOI of 1 with the respective virus at 37°C for 16h. On the following day, antibody dilutions were prepared using a start concentration of 100 μg/ml. The antibodies were serially diluted 1:3 and then added in duplicates to the cells. The human derived monoclonal antibody CR9114 (42) was included as a positive and an irrelevant anti-Lassa virus antibody (KL-AV-1A12 (45)) was used as a negative control. Next, 75,000 effector cells/well were added to the plate and incubated for 6 h at 37°C.The luciferase substrate was added in the dark and the luminescence activity measured after 5 minutes using a Synergy Hybrid Reader (BioTek). The data was analysed using GraphPad prism 7.

### NA inhibition assay

NA inhibition assays were performed as described in detail earlier (7, 54). In brief, 96-well flat bottom Immulon 4 HBX plates (ThermoFisher Scientific) were coated with 150 μl/well of fetuin (Sigma Aldrich) at a concentration of 50 μg/ml overnight at 4°C. The following day, the N6 mAbs were serially diluted 1:3 in PBS with a starting concentration of 30 μg/ml. The respective viruses were diluted in PBS at 2x the 50% effective concentration (EC50), which was determined in an NA-assay previously. The virus (75 μl/well) was then added to the mAb dilution plate and the virus/mAb mixture was incubated for 1h 45 min shaking at RT. During this time, the fetuin coated plates were washed 6x with PBST and then blocked with 5% BSA/PBS for at least 1h at RT. After 1 hour, the fetuin plates were washed and 100 μl of the virus/antibody mixture were added to plates and incubated for 2h at 37°C. The plates were washed and incubated with 5μg/100μl/well of PNA for 1h 45 min in the dark. The plates were then washed and developed by using 100 μl/well Sigmafast OPD solution (Sigma Aldrich). After 7 minutes of incubation in the dark, the reaction was stopped by adding 50 μl/well of 3M HCl and the plates read at an absorbance of 490 nm using a Synergy H1 hybrid multimode microplate reader (BioTek). The IC50 values were calculated by using GraphPad Prism 7.

### Plaque reduction neutralization assay

Plaque reduction neutralization assays were performed on MDCK cells (300,000 cells/well) which were seeded in a sterile 12-well plate the day before. The following day, the mAbs were serially diluted 1:3 in 1x MEM and 50 μl of A/duck/Czechoslovakia/1956 (H4N6, 1000 PFU/ml) were added to each dilution and incubated shaking at RT for 1h. The cells were washed with PBS and the antibody-virus mixture was added for 1h at 37°C. The mixture was then aspirated and the cells overlaid with agar consisting out of minimal essential medium (2xMEM), 2% Oxoid agar, 1% DEAE, TPCK-treated trypsin as well as the respective antibody. The plates were incubated at 37°C for 2 days and the cells afterwards fixed with 3.7% PFA in PBS. The plaques were then visualized by immunostaining. Briefly, the agar-overlay was removed and the plates blocked with 3% milk/PBS for 1h at RT. Afterwards, an antibody-cocktail made out of all ten N6-antibodies diluted 1:3000 in milk/PBS was added and incubated for 2h at RT. The plates were washed and incubated with secondary antibody (Antimouse IgG (H&L) Antibody Peroxidase Conjugated, Rockland) diluted 1:3000 in 1% milk/PBS for 1 h. The plates were washed and developed by using Trueblue reagent (KPL). The plaque number as well as the plaque size was determined for each antibody dilution and the percent inhibition determined based on a no-antibody control. The data was analysed using GraphPhad Prism 7 (54, 55).

### Evaluation of the prophylactic efficacy in mice

To test the prophylactic efficacy of the N6 antibodies, female 6-8 week old DBA/2J mice (The Jackson Laboratory) received an antibody dose of 5mg/kg intraperitoneal (n=5 per group). An irrelevant anti-Lassa antibody (KL-AV-1A12 (45)) was given to one of the groups as negative control. Two hours after antibody administration, the mice were anesthetized and intranasally challenged with 5x mLD_50_ (mouse lethal dose, 50%) of A/duck/Czechoslovakia/1956 (H4N6) or A/Shenzhen/1/2016 (H5N6). Weight loss was monitored over 14 days and any mouse which lost more than 25% of its initial body weight was sacrificed. Survival and weight loss data were analysed by using GraphPad Prism 7. All animal procedures were performed in accordance with the Icahn School of Medicine at Mount Sinai Institutional Animal Care and Use Committee (IACUC).

### Escape mutants

MDCKs were plated at a density of 1,000,000 cells/well in a 6-well plate the day prior The next day, cells were washed with PBS and infected with an MOI of 1 with the virus A/duck/Czechoslovakia/1956 (H4N6) by using 1xMEM as well as 1 ug/ml TPCK-treated trypsin. The respective mAbs were added at a concentration of 1×IC_50_ to the virus mixture and the cells then incubated for 2 days at 37°C. The cell supernatant was collected and used to continue the passaging of the virus. Additionally, to this, the IC50 concentration of the mAbs was doubled with every passage. This procedure was repeated until an IC50 concentration of 128 was reached. The presence of the virus within the samples was confirmed by performing an immunostaining of the cell layer after every passage. The same virus was passaged in parallel completely without mAbs as well as together with an irrelevant antibody (anti-Lassa KL-AV-1A12 (45)), to confirm that no random mutations were acquired over time.

Once the last passage was reached, the viruses were plaque purified and injected, together with the respective mAb, in 10-day old embryonated chicken eggs, to propagate the virus. The eggs were incubated for 2 days at 37°C. RNA was extracted from the allantoic fluid by using the Direct-zol RNA isolation kit from Zymo research. The purified RNA was used to deep-sequence the escape viruses.

## Acknowledgements

This work was supported in part by the National Institute of Allergy and Infectious Disease (NIAID) Centers of Excellence for Influenza Research and Surveillance (CEIRS) contract (HHSN272201400008C).

